# Altered inflammatory state and mitochondrial function identified by transcriptomics in paediatric congenital heart patients prior to surgical repair

**DOI:** 10.1101/2024.02.05.577755

**Authors:** Francesca Bartoli-Leonard, Amy G. Harris, Kelly Saunders, Julie Madden, Carrie Cherrington, Karen Sheehan, Mai Baquedano, Giulia Parolari, Andrew Bamber, Massimo Caputo

**Affiliations:** Bristol Medical School, Faculty of Health Sciences, University of Bristol, Bristol, UK; Bristol Heart Institute, University Hospital Bristol and Weston NHS Foundation Trust, Bristol, UK; North Bristol NHS Trust, Westbury on Trym, Bristol, UK

## Abstract

**Objective:** Congenital heart disease (CHD) remains the most common birth defect, with surgical intervention required in complex cases. Right ventricle (RV) function is known to be a major predictor in sustained cardiac health in these patients, thus by elucidating divergent profiles between CHD and control through tissue analysis this study aims to identify new avenues of investigation into the mechanisms surrounding reduced RV function.

**Approach & Results:** Transcriptomic profiling, *in silico* cellular deconvolution and functional network analysis was conducted on RV biopsies obtained from CHD and control paedatric patients. Analysis identified an increase in mitochondrial dysfunction genes *RPPH1* and *RMPR* (padj = 4.67^E-132^, 2.23^E-107^, respectively), Cytotoxic T cell markers *CD8a, LAGE3* and *CD49a* (p = 0.0006, p < 0.0001, p = 0.0118, respectively) and proinflammatory marker *Caspase1* (p=0.0055) in CHD compared to control. Gene set enrichment identified mitochondrial dysfunctional pathways, predominately changes to the oxidative phosphorylation processes. Negative regulation of mitochondrial functions and metabolism was identified in functional network analysis, with dysregulation of mitochondrial complex formation. Histological analysis confirmed an increase in cellular bodies with the CHD RV tissue, and positive staining for both CD45 and CD8 in CHD RV tissue, which was absent in control. Deconvolution of bulk RNAseq data suggests a reduction in CD4+ T cells (p = 0.0067) and an increase in CD8+ T cells (p = 0.0223). Network analysis identified positive regulation of the immune system and cytokine signalling clusters within the inflammation functional network as were lymphocyte activation and leukocyte differentiation.

**Conclusions:** Utilizing RV tissue from paediatric patients undergoing CHD cardiac surgery this study identifies dysfunctional mitochondrial pathways and an increase in inflammatory T cell presence prior to reparative surgery.

## Introduction

Congenital heart disease (CHD) remains the most common birth defect, with surgical intervention required every 5 years in children^1^ to every 15 years in adults^2^, conferring significant risk of mortality with each reintervention. In contrast to adult heart disease, which disproportionately affects the left ventricle, complex CHD can be characterized by increased right ventricular (RV) pressure, leading to RV hypertrophy and eventually failure, with RV function known to be a major predictor in survival^3,4^.

Current treatment strategies for left ventricular dysfunction such as β-blockers, iACE and Angiotensin II receptor blockers fail to impact RV dysfunction in CHD patients. This lack of viable treatment is further compounded by the current available diagnostic modalities failing to predict early-stage subclinical changes occurring in the RV, highlighting the knowledge gap within this pathology. Alongside RV dysfunction in CHD, low level inflammation in the myocardial tissue within malformed hearts promotes further dysfunction over time^5^. Small animal studies have demonstrated that non-ischemia heart failure and RV failure is orchestrated by the adaptive immune response^6,7^, with an exhausted T cell population identified in RV dysfunction at a single cell level^8^. Understanding the inherent differences in RV tissue from CHD patients compared to controls may suggest new ways to adapt surgical treatments to improve outcomes in patients, and moreover may highlight the critical yet subtle changes to pay attention for in post-surgical patients. Thus, this study hypothesized that activated inflammatory pathways and mitochondrial dysfunction may be present in the RV prior to and during surgical intervention, leading to the increase in inflammatory cell prevalence within the RV. Improved understanding of the complex mechanisms occurring within the RV within CHD patients may aid the development of therapeutics to mitigate mitochondrial dysfunction and improve RV outcomes post-surgery in this high-risk paediatric cohort.

## Methods

### Study Population

Twenty-four RV samples were collected from discarded myocardial tissue from patients undergoing cardiac surgery at Bristol Royal Hospital for Children, following informed consent or from control autopsy subjects (CHD RNAseq; 14, CHD histology; 5, control histology; 5). All samples were taken from patients between the age of 0 and 16 years old. Of the CHD samples, all were diagnosed with a cyanotic pathology shortly after birth; 11 Tetralogy of Fallot (ToF), two Truncus Arteriosus and one critical pulmonary stenosis. Cardiac tissue was obtained intraoperatively and submerged immediately in liquid nitrogen, stored at -80°C and processed for downstream use. All patients underwent cardio-pulmonary bypass. Control tissue was obtained wax-embedded from clinical pathology. This study was approved by NHS REC 19/SW/0113 in accordance with the declaration of Helsinki. Clinical data, pre- and post-operative echocardiography and clinical diagnosis were reviewed for all patients retrospectively to confirm diagnosis. Patients with suspected or diagnosed genetic pathologies were excluded.

### Tissue processing and Histological Analysis

Fresh biopsies (10 mg net weight) were collected from surgery from the apex of the RV immediately after institution of cardiac pulmonary bypass and snap frozen upon harvesting or processed immediately for wax embedding (Fig 1A). Briefly, RV tissue was embedded in paraffin and 6 μm sections were cut using a microtome (Leica, Germany) and slides dried in a heated chamber. Sections were deparaffinized in xylene and decreasing ethanol concentrations as previously described^9^. Slides were then washed in PBS for 5 minutes, before incubating in heated sodium citrate for 30 minutes and blocking in 0.3% H_2_O_2_ in PBS for 20 minutes. Slides were then washed in tap water and placed in PBS for 5 minutes. Next, slides were blocked in 4% bovine serum albumin (Sigma, UK) at room temperature for 1h in a humidified chamber. Following blocking, slides were incubated with primary antibodies (Supplementary Table 1) diluted in PBS with 2% BSA at room temperature for 1 hour in a humidified chamber. Slides then washed three times in PBS for 5 minutes before diluted secondary antibodies (Supplementary Table 1) were added in PBS with 2% BSA at room temperature for 1 hour in a humidified chamber. Slides were then washed three times and incubated with streptavidin-HRP (Sigma, UK) at room temperature for 30 minutes before washing a further three times in PBS. Finally, slides were developed in DAB (Vector Laboratories, UK) and washed in tap water. Slides were then counterstained in Gils haematoxylin for 30 seconds before washing twice in tap water, placed in ammonium water for 15 seconds and then washed in water twice again. Slides were dehydrated as before and mounted using a DPX based mounting media and imaged on a Slideview VS200 (Olympus, USA).

**Figure 1.**
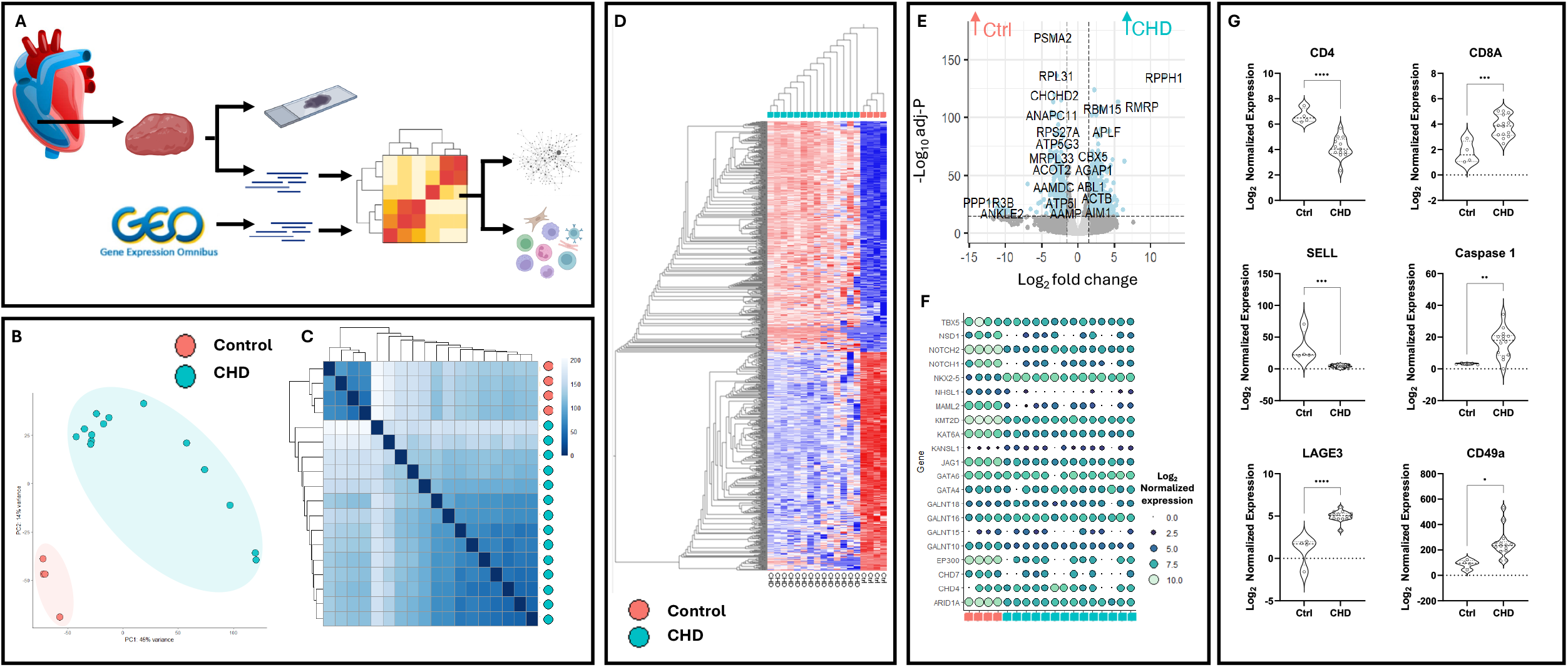
Transcriptomic analysis demonstrates a reduction in homeostatic genes and an increase in immune-infiltration regulators within congenital heart disease and control patients. Schematic workflow of sample processing and analysis. B) Principal component analysis of n=14 whole tissue right ventricle transcriptomic data & n=4 control tissue from paediatric patients, no cut off applied. C) Heatmap clustering of samples based on distance matrix with hierarchical clustering applied on normalized log transformed counts. D) Hierarchical clustering of log-transformed transcriptomic profiles with a cut-off of padj ≥ 0.05, logFC ≤ 2. E) Volcano plot with differentially expressed genes in control and CHD transcriptomics. F) Dot plot examining hallmark CHD genes, through log-normalized gene expression between control and CHD. G) Immune cell genes CD4, CD8A, T cell maturation marker *SELL*, inflammatory markers *Caspase 1, LAGE3* and T cell homing marker *CD49a*. Statistical analyses were performed using paired-T tests corrected for multiple comparisons (n=18). ^*^p < 0.05, ^**^ p < 0.01 ^***^p < 0.001.

Haematoxylin and Eosin staining was conducted on deparaffinized sections. Briefly, slides were washed in distilled water, incubated in haematoxylin for 30 seconds, rinsed in running tap water and Scotts’ tap water before incubated in eosin for 2 minutes. Slides were washed in ammonium water for 30 seconds before dehydrating and mounting as before and imaged on the Slideview VS200 (Olympus, USA).

Picro-Sirius Red was used to determine collagen fibres under polarized light. Briefly, deparaffinized slides were washed in distilled water and incubated in Picro-Sirius Red (Abcam, USA) for 60 minutes. Slides were then rinsed in Acetic acid solution (0.5%) until excess stain was removed. Finally, slides were dehydrated and cleared, before mounting with a DPX mounting media as before. Slides were imaged by polarized light and brightfield light microscopy and merged on the Slideview VS200 (Olympus, USA).

### Transcriptomic Analysis

Strand-specific bulk RNA sequencing was performed on whole tissue snap frozen RV biopsies via Genewiz (Azenta). Briefly, total RNA from RV was isolated from 10mg of tissue, with poly-A selection. Of the 24 samples submitted, 14 passed quality control and were taken forward for sequencing. Sequencing was conducted on Illumina NovaSeq, yielding 2x150bp paired end reads, 30 million read pairs per sample, with >80% of bases with Q30 or higher. Control samples were downloaded through the SRA toolkit, series GSE217772 containing raw RNA-sequencing FASTQ files of RV samples of unaffected paediatric patients (1-6 years old), who have undergone traumatic brain injury but maintained cardiac function. RVs were isolated shortly after death and snap frozen before processing and sequencing at the Children’s Hospital of Chongqing Medical University. Whole transcriptome was sequenced on an Illumina NovaSeq 6000 with paired end reads produced. FastQC (Babraham Bioinformatics, V0.12) was used to provide quality control on raw counts. Paired-end data was trimmed via SLIDINGWINDOW:4:20 command with Trimmomatic-0.35^10^. Quantification was conducted using Salmon^11^ (V1.10.2) via mapping-based mode *salmon -quant* indexed against human genome assembly GRCh38.

### Differential Enrichment and Network Analysis

DESeq2 (v1.40.2, R)^12^ was used to perform differential expression analysis. Briefly, tximportData was used to import salmon output files, stratified by the presence of CHD (CHD; 14, ctrl; 4) and results produced with the *apeglm* log2 fold change shrink. Principal component analysis was conducted with the *PlotPCA* function on normalized transformed data through the *normTransform* function, variance stabilizing transformation was used to log transform data. Gene clustering was performed with the *genefilter* package, with the top variable genes stratified by the presence of CHD through the transformed values of the matrix of normalized values, with volcano plots produced by *EnhancedVolcano*. Gene set enrichment was performed via *fgsea* R package, utilizing the KEGG, WikiPathways, Hallmark and Reactome databases and plotted via *ggplot2*. Network analysis was conducted inCytoscape. ClueGO Cytoscape plugin was used to visualize the non-redundant biological terms for large clusters, based on the databases above on the significantly upregulated genes within the CHD population. Network was based on the kappa statistics and reflects the relationship between the terms, based on the similarity of their functional genes.

### Transcriptomic Deconvolution

Bulk transcriptomic data was deconvoluted via *granulator* using the single cell matrix from healthy RV dataset GSE145154^8^. Raw data was downloaded through the SRA toolkit, and Seurat object produced as previously described^13^. Seurat object data was normalized with *scTRANSFORM* and celltype identified as dictated by the published data. Deconvolution was modelled via seven methods; *dtangle, nnls, ols, qprog, qprogwc, rls* and *svr*. Pearsons’ correlation of cell type proportions across all methods was assessed and *dtangle* was taken forward. Estimated cell type proportions were normalized to 100% and presented per donor.

### Statistical Analysis

Student t-test were used for 2-group comparisons of normally distributed data. ANOVA was used for >2 group comparisons, performed on continuous, normally distributed data. Bonferroni correction for multiple comparison test was applied where required.

### Data availability

Raw and processed transcriptomic data available through GEO (). All scripts used within this study can be made available upon reasonable request to the authors.

## Results

Transcriptomic analysis was conducted on 14 patients and 4 controls, to examine the transcriptomic profile of biopsies taken from RV during cardiothoracic surgery and assess the molecular landscape. Principal component analysis of RV transcriptomic profiles demonstrated clear clustering based on disease state with no cut off applied (Fig 1B). Distance matrix of log-normalized transcript counts presented as a hierarchical heatmap clustered control and CHD separately (Fig 1C), with no filtering applied.

Following DESeq2^12^ analysis, hierarchical clustering demonstrates the differences between control and CHD tissue (false discovery rate <0.05, abs[log fold-change]>2), with little variation seen in the global transcriptomic profile between CHD patients (Fig 1D). Mitochondrial dysfunction genes *RPPH1* (log2FC = 11.86, padj = 4.67 E-132) and *RMPR* (log2FC = 8.80, padj = 2.23 E-107) were significantly increased in CHD compared to control, as was the proinflammatory mediator *ABL1* (log2FC = 1.79, padj = 2.60 E-39). Mitochondrial regulator *CHCHD2* (log2FC = 3.89, padj = 8.38 E-117) was upregulated in control conditions alongside ribosomal subunits *PSMA2* (log2FC = 3.39, padj = 7.07 E-166) and *PRL31* (log2FC = 2.96, padj = 3.10 E-133) (Fig 1E). Expression of hallmark CHD genes was assessed in RV compared to control, with CHD RV samples showing a reduction in *TBX5*, JAG1, *NOTCH2* and *NOTCH1* (padj = 2.24E-07, 1.38E-27, 7.76E-31 and 9.11E-7 respectively). Notably, no change was detected in *GATA4* or *GATA6* genes directly associated with CHD development^14^ (padj = 0.79, 0.80 respectively), however the *NKX2-5*, linked to 10 CHD cyanotic abnormalities was increased in CHD compared to control, highlighting the complex genetic variation within these patients (padj = 4.97E-16) (Fig 1F). Following previous studies of dysregulated immune responses within CHD patients, key adaptive immune cell markers were assessed within the RV. T helper cell marker *CD4* was significantly downregulated (p < 0.0001), as was T cell naivety marker *SELL* (p = 0.003). Cytotoxic T cell marker *CD8a* was significantly increased in CHD compared to control (p = 0.0006), as was proinflammatory marker *Caspase1* (p=0.0055). CD8+ T cell accumulation marker *LAGE3* and T cell tissue localization marker *CD49a* were both increased in CHD compared to control (p < 0.0001, p < 0.0118, respectively) (Fig. 1G), of note, no individual B cell markers were identified within the transcripts.

Deconvolution of the transcriptomic profiles was conducted to assess the cellular composition of the RV biopsies, utilizing a single cell backbone of RV biopsies in dilated cardiomyopathy and healthy control taken from adult donors^8^ to predict cellular proportions. As expected, transcripts clustered as B cells had a unique transcriptome compared to other cell types, with cardiomyocytes and fibroblasts clustered more closely. T cells, NK cells, myeloid cells and endothelial cells clustered similarly (Fig 2A). Normalized proportions of each cell type (Fig 2B) demonstrated similarity between disease and control, irrespective of specific CHD malformation. Estimated proportions of cell types within both control and CHD (Fig 2C) predicted cardiomyocytes made up the majority of the cells present within the RV (48% ± 7.42, and 70% ± 16.41, respectively, p < 0.001). No difference was observed with the prevalence of fibroblasts (6.7% ± 1.51 and 6.2% ± 7.94, p = 0.4236) with the endothelial cell fraction exhibiting significantly more within the control compared to the CHD (22% ±3.63, 9% ±4.49, p = 0.0002). No difference in B cells was observed (1% ± 0.56, 2% ±0.90 p = 0.235), with a decrease observed in natural killer (3% ± 2.58, 7% ± 2.11, p = 0.0043), myeloid (2% ± 1.07, 5% 1.16, p = 0.0001) and CD4+ T cells (2% ± 1.90, 6% ±2.76, p = 0.0067) in CHD compared to control. Notably, CD8+ T cell prevalence was greater in the CHD population compared to control (7% ±4.7, 4% ± 2.9 p=0.0223) (Fig 2D), with no association between CHD pathology and CD8+ prevalence. Histological analysis of RV biopsies obtained from the same patients with CHD (n=5) and novel controls (n=5) obtained from Bristol pathology department (unique from the RNAseq samples) showed an increase in cellular bodies within CHD RV tissue compared to control. Breakdown of type I collagen in line with increased RV fibrosis was observed in CHD compared to control, demonstrated by a lack of red/yellow fibres shown in Picro-Sirus red staining (PSR). Immuno-histological staining of immune markers CD45 and CD8 were positive in CHD and respectively less stained in control tissue (Fig 2E).

**Figure 2.**
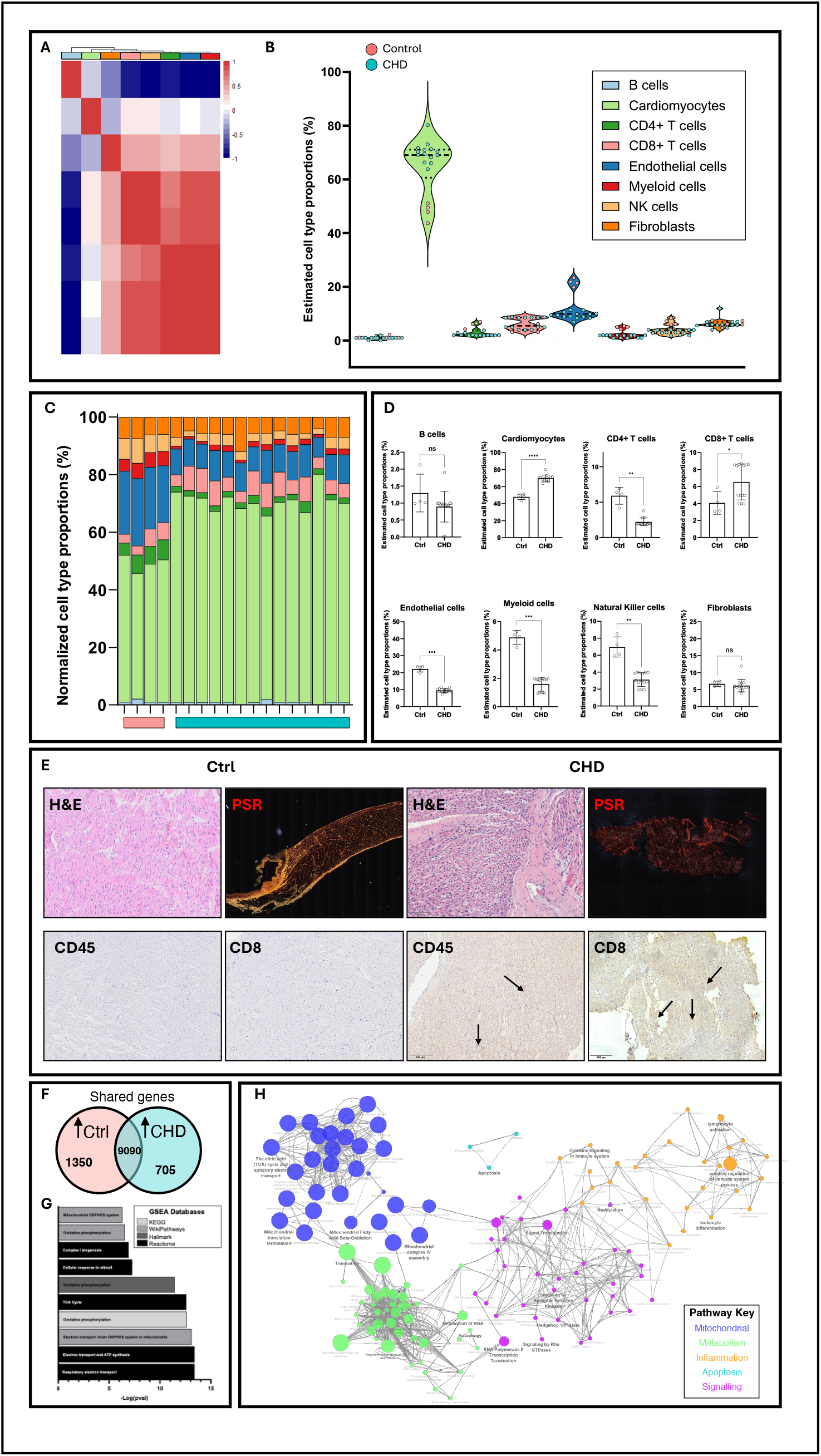
Deconvolution of transcriptomic profiles and network analysis of get set enrichment identifies a dysregulation in inflammatory and mitochondrial pathways. A) Correlation heatmap of cellular phenotypes deconvoluted from RNAseq data using healthy and inflamed right ventricle single cell RNAseq datasets in human heart failure. B) Estimated cell type proportions from deconvoluted RNAseq datasets based on log-normalized TPM data. C) Estimated cell type proportion for each donor, normalized to 100%. D) Cell types identified in deconvolution. E) Histological analysis; hematoxylin & eosin, Picro-Sirus red, CD45 and CD8 of control vs CHD RV biopsies. F) Whole transcriptomic analysis following alignment, 82% of genes identified overlap. G) Top 15 significant (p < 0.05) gene set enrichment pathways in CHD identified from 5 databases; humanPhenotype Ontology, KEGG, WikiPathways, Hallmark gene set and Reactome. H) Network visualization of genes increased in CHD vs control tissue. Size corresponds to inverse Bonferroni corrected pval; largest size; smallest pval.

Transcriptomic analysis identified 10440 gene transcripts within control and 9795 within CHD with 82% shared homology (Fig 2F). Gene set enrichment analysis of differentially expressed genes, upregulated within the CHD RV biopsies identified the top 15 pathways from HumanPhenptype Ontology, KEGG, WikiPathways, Hallmark and Reactome databases, accessed through the *fgsea* R package. Mitochondrial pathways were predominantly enriched in the CHD samples, with abnormalities in the mitochondrion and changes to the mitochondrial respiratory electron transport chain (OXPHOS) reported (Fig 2G). Functional network analysis visualized non-redundant biological terms for large gene clusters within the transcriptomic profile of CHD RV biopsies (Fig 2H). Five main clusters were identified pertaining to mitochondrial functions, metabolism, inflammation, apoptosis and signalling pathways. Mitochondrial related functional clusters were the most significantly upregulated within CHD, which was closely linked to dysregulated metabolic functions. Positive regulation of the immune system and cytokine signalling clusters were increased within the inflammation functional network, as well as lymphocyte activation and leukocyte differentiation, giving rationale to further assess the immune cell compartment and metabolic function within the right ventricles.

## Discussion

Surgical intervention for CHD is currently the only form of treatment, however the majority of these patients require multiple reinterventions throughout their life, with fibrosis, occlusion and calcification limiting the lifespan of implanted materials^15-17^. Animal models, predominantly murine and porcine recapitulate the surgical aspect of CHD, however, due to the abrupt change from *healthy* to pathogenic through intervention, they fail to recapitulate the developmental aetiology of CHD and the slow changes in pressure and cellular composition. Thus, this study set out to profile tissue taken directly from patients immediately prior to surgery to better understand the differential cellular landscape within the right ventricle as a means to identify potential targetable differences between disease and control. By transposing RNAseq data onto a single cell backbone, cellular ratios can be estimated to elucidate inflammatory cells potentially active within RV tissue prior to intervention, which may impact the response of the localised tissue to implantable grafts, leading to premature failure and the need for repeated interventions. Landmark studies have highlighted the increased pro-inflammatory markers present within the myocardium of CHD patients^18,19^, with an altered immune profile reported in patients who underwent thymectomies routinely as part of cardiothoracic surgery^20^, exhibiting an increased level of T cell expression. The data presented here demonstrates a CD8+ T cell rich RV environment, identified through CD8+ T cell accumulation and infiltration marker LAGE3, tissue residency marker CD49a+, pro-inflammatory cytokine caspase-1 RNA expression and positive CD8+ and CD45+ histological staining. Previous studies have identified an increase in pro-inflammatory cells within adult heart failure patients^8^ and genetic malformations such as DiGeorge syndrome^21^, but have been thus far unconfirmed in non-genetically influenced CHD pathologies alone. Due to the clearly difficult nature of accessing tissue biopsies prior to surgery to assess risk, identification of circulating markers such as increased inflammatory cell populations or their subsequent cytokine profile may be able to act as surrogate to better understand risk for these patients pre-surgery, though further targeted studies must be taken to fully elucidate this tentative relationship. Within adult populations, pre-operative inflammatory status has been demonstrated to impact surgical cardiac outcomes^22^, with a systemic inflammatory index^23^, currently utilized in cancers and stroke able to accurately predict outcome favourability, however the efficacy of using such scoring in a paediatric population remains unfounded.

Network analysis conducted on the differentially expressed RNA profiles in CHD confirmed an inflammatory profile and moreover identified a network of mitochondrial dysfunction associated genes, suggesting a change in RV metabolic profile prior to surgical intervention. Mitochondrial dysfunction defines T cell exhaustion^24,25^, and accumulating evidence has suggested that mitochondrial dysfunction and low-grade inflammation plays a critical role in both the development of RV maladaptation^26^, the progression of inflammation and accumulation of inflammatory cells within regions of dysfunction^27,28^. Depletion of mitochondrial DNA and impaired mitochondrial replication have been associated with early events of RV hypertrophy, preceding clinical diagnosis^29^. In reperfusion injury following cardiac surgery procedures, ischemic or reperfusion injuries to the myocardium can significantly damage the mitochondrial structure and function^30,31^ and thus if the mitochondria is already damaged or dysfunctional in CHD patients prior to surgery, outcomes may be significantly worse. Metabolic switching from glucose oxidation to glycolysis due to pre-clinical ischemia, may present an opportunity for a therapeutic intervention. Pre-surgical treatment to promote restorative and *healthy* mitochondrial function within the right ventricle, such as targeted pyruvate dehydrogenase kinase inhibitors^32^ and fatty acid oxidation inhibitors^33^ may promote enhanced recovery and longer-term stability of the RV post-surgery. However, further mitochondrial functionality assessments and effect on the left ventricle must be conducted on fresh tissue to target the mitochondrial dysfunction further.

In summary, the work presented here identifies dysfunctional pathways present within RV biopsies from patients with CHD. Whilst animal models can attempt to recapitulate such aetiologies, to understand the complex mechanisms at play which influence the outcomes of surgical intervention in these patients, more studies must be undertaken on primary tissue, integrating novel ways to utilize pre-existing -*omic* libraries. Due to the fragile nature of such patients, further non-invasive studies must be conducted to broaden the understanding of the complex mechanisms within paediatric CHD.

## Acknowledgments

Thomas Mitchell of University Hospital Bristol and Weston NHS Foundation Trust for proof reading this manuscript.

